# A synthetic stroma-free germinal center niche using material-surface driven polyvalent signaling efficiently induces antigen-specific humoral immunity *ex vivo*

**DOI:** 10.1101/194456

**Authors:** Kyung-Ho Roh, Hannah K. Wilson, Pallab Pradhan, Kevin Bai, Caitlin D. Bohannon, Gordon Dale, Jardin Leleux, Joshy Jacob, Krishnendu Roy

## Abstract

B cells play a major role in the adaptive immune response by producing antigen-specific antibodies against pathogens and imparting immunological memory. Following infection or vaccination, antibody-secreting B cells and memory B cells are generated in specialized regions of lymph nodes and spleens, called germinal centers. Here, we report a fully synthetic *ex-vivo* system that recapitulates the generation of antigen-specific germinal-center (GC) like B cells using material-surface driven polyvalent signaling. This synthetic germinal center (sGC) reaction was effectively induced using biomaterial-based artificial “follicular T helper cells (T_FH_)” that provided both natural CD40-CD40L ligation as well as crosslinking of CD40; and by mimicking artificial “follicular dendritic cells (FDC)” to provide efficient, polyvalent antigen presentation. The artificial sGC reaction resulted in efficient B cell expansion, immunoglobulin (Ig) class switching, and expression of germinal center phenotypes. Antigen presentation during sGC reaction selectively enhanced the antigen-specific B cell population and induced somatic hyper-mutations for potential affinity maturation. The resulting B cell population consisted primarily of GC-like B cells (centrocytes) as well as some plasma-like B cells expressing CD138. With concurrent cell sorting, we successfully created highly enriched populations of antigen-specific B cells. Adoptive transfer of these GC-like B cells into non-irradiated isogeneic or non-lethally irradiated congenic recipient mice showed successful engraftment and survival of the donor cells for the 4 week test period. We show that this material-surface driven sGC reaction can be successfully applied to not only splenic B cells but also B cells isolated from more therapeutically relevant sources such as peripheral blood mononuclear cells (PBMCs), thus making our current work an exciting prospect in the new era of personalized medicine and custom-immunotherapy.

## Introduction

As a critical arm of adaptive immunity, B cells provide humoral immunity by producing antibodies towards various pathogens. In particular, the generation of memory B cells and long-lived plasma cells (LLPCs) is critical to immune memory and protection from recurring infection. Memory B cells recognize and are rapidly activated by recurring antigens to secrete antigen-specific high-affinity antibodies. LLPCs produce and maintain the protective level of these antibodies in the serum.

Memory B cells and LLPCs are typically generated inside special sub-anatomical microenvironments called germinal centers (GCs) within secondary lymphoid organs such as lymph nodes and spleen. In these lymphoid tissues, a small fraction of naïve B cells recognize a given antigen presented by follicular dendritic cells (FDCs); they then process and present the antigen to follicular helper T (T_FH_) cells. B cells forming close interactions with T_FH_ cells are further activated by costimulatory signals provided by T_FH_ cells, and these activated B cells proliferate rapidly, expanding populations of responding B cell clones within the GC. Mimicking this complex set of events *ex vivo* has proven elusive. Such *ex-vivo* synthetic GCs (sGC) could not only provide new insights in B cell biology, but also allow us to generate antigen-specific B cells for antibody production and adoptive immunotherapies.

We hypothesized that we can recapitulate the GC reaction *ex vivo* by effectively synthesizing the critical signaling events provided by T_FH_ cells and FDCs (**Fig. 1a**) in the presence of an antigen. Traditionally, *in-vitro* B cell activation has been achieved by engagement of CD40 molecules on B cell membrane using anti-CD40 antibodies^1^, recombinant (CD40L)^2-4^, or co-culture with supporting cell lines that have been genetically modified to express CD40L^5,6^. Recently, Kitamura and colleagues have developed a fibroblast-based feeder cell line that expresses both CD40L and B-cell activating factor (BAFF), particularly for the purpose of generating *ex-vivo* GC reactions^7,8^. It was also reported that culturing these feeder cells in a hydrogel composed of Arg-Gly-Asp(RGD)-presenting extracellular matrix enhanced the co-cultured B cell survival and differentiation^9,10^. Despite these previous efforts, it has not been explored whether a purely synthetic, biomaterials-based design can effectively recapitulate CD40L presentation by T_FH_ cells and antigen-presentation by FDCs in the context of modulation of GC reaction and generation of antigen-specific, functional B cells. Enabling mimicry of GC reactions *ex vivo* without the use of genetically-modified feeder cell lines offers significant advancement in designing potential applications of sGC reaction such as i) providing a simpler, cheaper, and more rapid model for studying GC B cell physiology and pathology of B cell malignancies, ii) development of multiplex screening platforms for drugs and immunotherapeutics, and iii) *ex-vivo* induction of effective humoral immunity as a potential B cell immunotherapy regimen, especially in immunocompromised patients.

**Figure 1.**
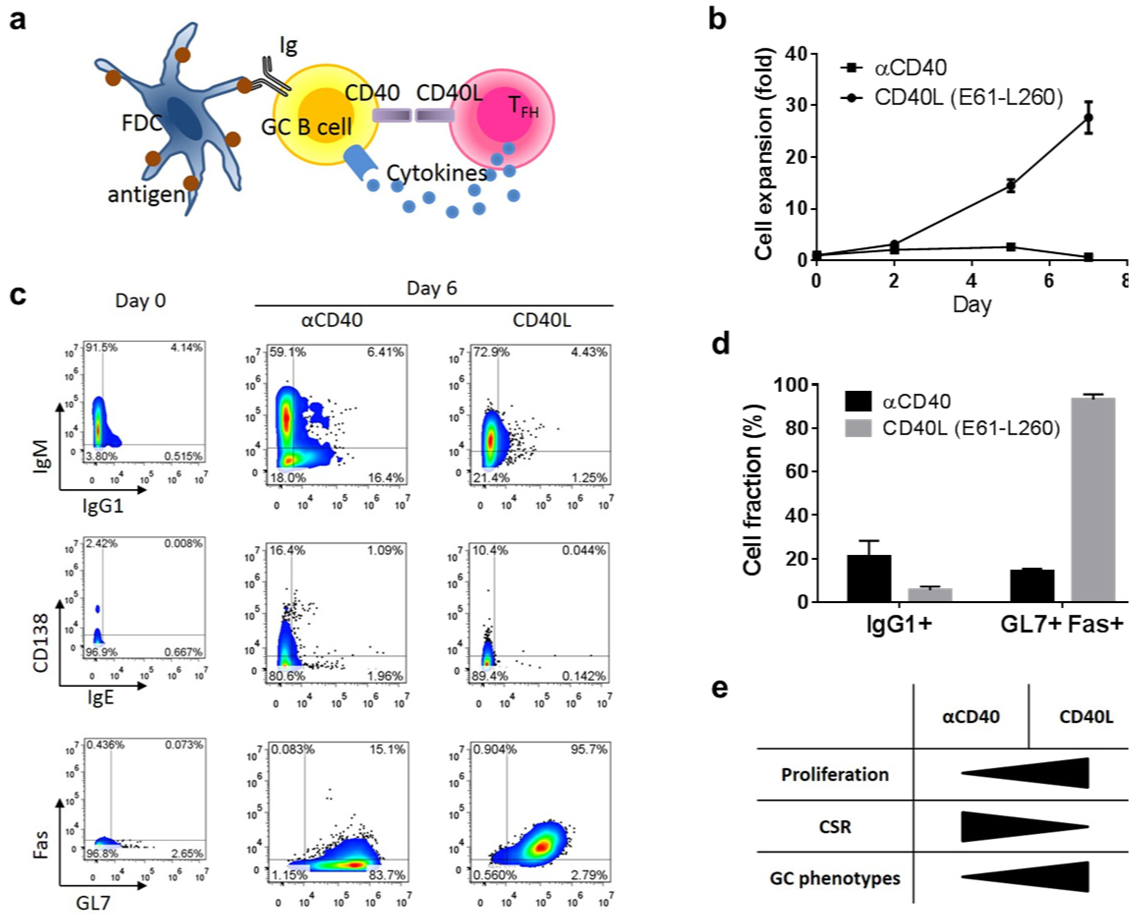
Incompetence of soluble anti-CD40 antibody and CD40L in induction of full B cell activation. (**a**) Schematic illustration of critical signaling components required for the B cell activation in a GC microenvironment. (**b**) Cumulative fold increase in the number of live B cells. The squares (αCD40) and circles (CD40L) are mean values of each time point and the error bars are s.d. from 3 experiments. (**c**) Representative sets of flow cytometry analysis for the indicated markers before (day 0) and after (day 6) of B cell cultures using soluble αCD40 or CD40L in conjunction with IL-4 and BAFF. Numbers indicate the percentages of cells in the respective quadrants. (**d**) Summary of flow cytometry analysis plotted for the percentages of cells in the indicated gates on day 6 with mean ± s.d. from three separate experiments. (**e**) Comparison of different quality and quantity that soluble αCD40 or CD40L induce on B cells.

The selective expansion of antigen-specific B cell clones is initiated by the effective antigen presentation by FDC. As these B cell clones proliferate in the GCs, the affinities toward antigenic epitopes are increased by somatic hypermutation (SHM) of immunoglobulin (Ig) antigen receptor genes under a concurrent selection mechanism^11-13^ for high-affinity clones (affinity maturation). There has been little effort to emulate this critical aspect of the GC reaction *ex vivo*. If successful, such sGC reaction and *ex-vivo* generation of high quality and high affinity antigen-specific B cells could provide significant advancement in treating various diseases that require antigen-specific humoral immunity.

Here, we report that GC reactions can be effectively recapitulated *ex vivo* by purely biomaterials-based artificial T_FH_ and FDCs with polyvalent presentation of ligands or antigens. By testing and comparing various CD40 ligation methods for their capability to induce expansion, class switch recombination, and expression of GC B cell-specific phenotypes, we found that surface-density controlled CD40L presentation by microbeads were the most effective as artificial T_FH_ cells. Additionally, microbead surface-bound or multimeric antigen presentation were successfully employed to create enriched populations of antigen-specific B cells that had undergone class-switched and somatic hypermutation. Importantly, these sGC reactions were applied to not only spleen-isolated B cells but also the B cells isolated from peripheral blood mononuclear cells (PBMCs), a critical aspect for clinical translation. *Ex-vivo* sGC-generated B cells engrafted to the spleen and the lymph nodes following adoptive transfer, suggesting that these cells can be potentially used in immunotherapy as a novel means to confer antigen-specific humoral immunity.

## Results

### Ligation of CD40 by either soluble anti-CD40 antibody or soluble recombinant CD40L (CD154) induces *incomplete* B cell activation and GC reaction

For effective activation of B cells in the GC reaction, both CD40-CD40L (CD154) interaction and the soluble factors secreted by T_FH_ and FDCs - including interleukins (e.g. IL-4) and B cell activating factor (BAFF) - provide critical signals^14^. (**Fig. 1a**). As soluble anti-CD40 antibody (αCD40) or recombinant CD40L have traditionally been used for B cell activation, we first tested the *quality* and *quantity* of B cell activation that these soluble CD40 ligations induce. In the presence of IL-4 and BAFF, splenic B cells were cultured with either αCD40 (Clone 1C10) or CD40L (E61-L260) for 7 days. In both cultures, B cells formed spheroid-looking aggregates within 2 days. But only the culture with CD40L yielded significant B cell expansion (**Fig. 1b**). For B cells activated by αCD40, cell viability measured on Day 6 was only about 2∼5%, thus implying that αCD40 culture induces apoptosis. Flow cytometry was employed to examine class switch recombination (CSR), differentiation into plasma cells, and expression of GC phenotypes (**Fig. 1c, d**). A moderate level of CSR from IgM to IgG1 and expression of CD138, the plasma cell marker, was achieved in the culture using αCD40 compared to the minimal CSR and CD138 expression in the culture with CD40L. Conversely, CD40L was superior in induction of GC phenotypes (GL7+ Fas+) to αCD40. Thus in the perspective of GC reaction, αCD40 was more effective in induction of CSR than CD40L, but CD40L was superior to αCD40 in B cell expansion and in differentiation to GC-like B cells (**Fig. 1e**). Therefore, we could conclude that both natural ligation of CD40 using CD40L and crosslinking of CD40 might be necessary to induce an effective synthetic GC (sGC) reaction.

### Microbead-based artificial T_FH_ design presenting polyvalent CD40L with a controlled surface density effectively induced sGC reaction

In order to induce effective crosslinking of CD40 while leveraging the natural CD40-CD40L ligation, we designed a microbead-based artificial T_FH_ cell using recombinant CD40L tagged with HA peptide and iron oxide microbeads coated with anti-HA antibodies (**Fig. 2a**). Using this artificial T_FH_, sGC reaction was induced in presence of BAFF and IL-4. Splenic B cells quickly formed a complex with these artificial T_FH_, and grew as an organoid structure mimicking physiological germinal centers within 48 hours (**Fig. 2b**). As these sGC structures developed, areas enriched with artificial T_FH_ microbeads (darker under phase contrast optical microscope) often separated from the areas enriched with B cells.

**Figure 2.**
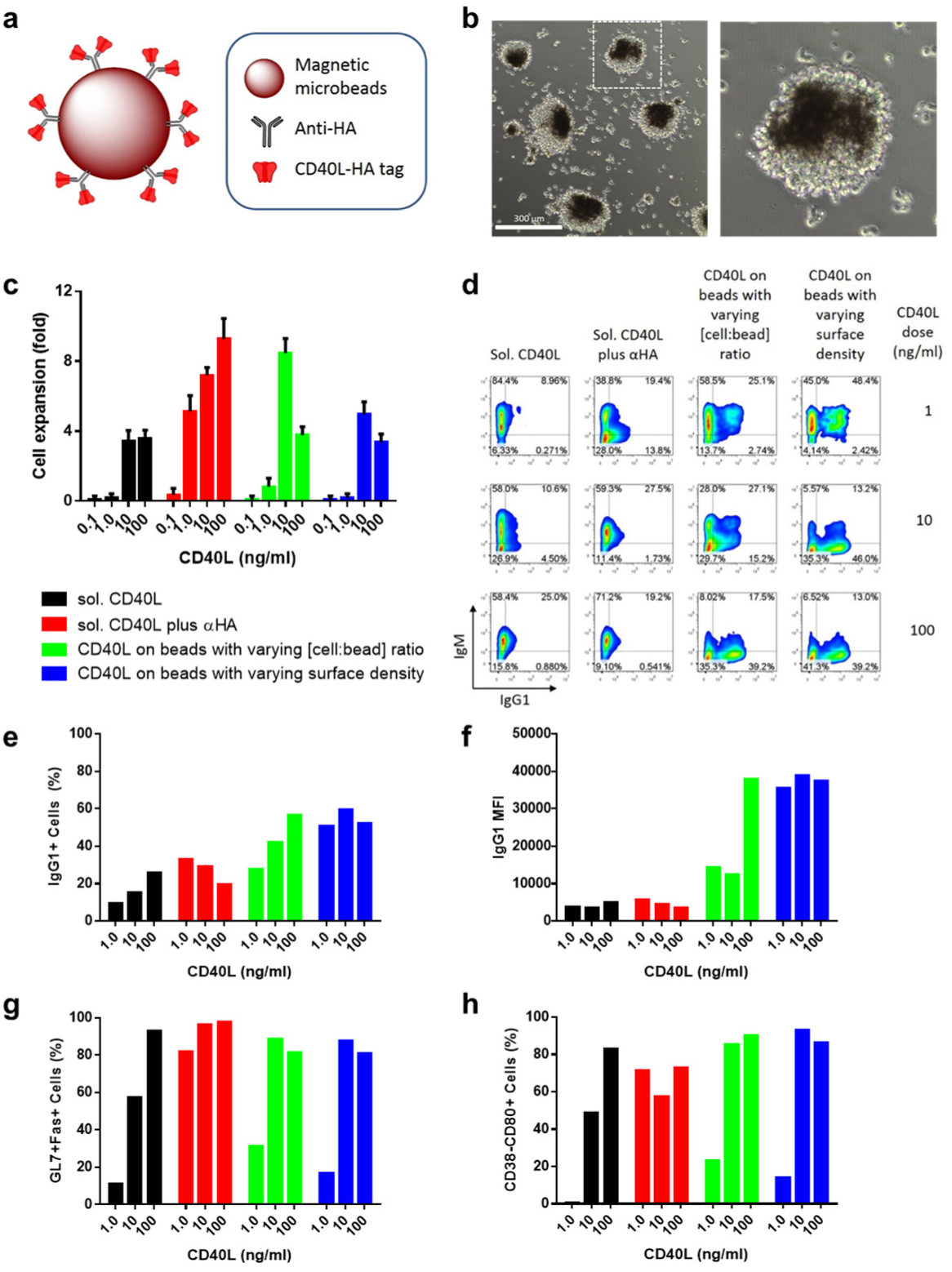
Artificial TFH design for induction of full activation of B cells and development of artificial GC structure and phenotypes. (**a**) Schematic representation of artificial TFH cell design. Surface density can be controlled simply by varying incubation concentration of CD40L molecules. (**b**) Phase-contrast light microscope images of artificial GC structures. Splenic naïve B cells complexed with artificial TFH microbeads develop into these spheroid structures within 48 hours of culture. It is often that darker area enriched with microbeads (artificial T cell zone) coexist with lighter artificial B cell zone. Scale bar (left) is 300 μm. Right image is an enlargement of dotted-line inset area of the left image. (**c-h**) Direct comparison of soluble CD40L and microbead-bound CD40L for activation of B cells and induction of GC reaction. For each indicated method of CD40-engagement method, the dose of CD40L was varied. (**c**) B cell fold expansion plot showing mean and s.d. of the number of live cells on day 6 of culture. Data was acquired from 5 wells per group from a single experiment. (**d-f**) Flow cytometry analysis for isotype class switching recombination (CSR) of B cells on day 6 of culture for each indicated condition. Percentage of IgG1+ B cell (**e**) and IgG1 MFI (**f**) were calculated and plotted from the representative flow cytometry chart (**d**). (**g, h**) Differentiation into GC-like B cells on day 6 of culture for each indicated condition. Percentage of B cells possessing two typical GC B cell phenotypes, GL7+Fas+ (**g**) and CD38-CD80+ (**h**), were calculated and plotted from respective flow cytometry analysis chart (**Supplementary Fig. 1**, **2**).

The unique advantage of such a biomaterial-based sGC design is that critical reaction parameters can be quantitatively controlled and tested. Specifically, we investigated how the sGC reaction can be modulated by controlling the dose of CD40L, the surface density of CD40L, and the ratio of B cells to artificial T_FH_ microbeads. sGC reaction conditions using soluble CD40L, with or without the soluble crosslinking antibody, were also tested for direct comparisons. First, the dose of CD40L was optimized by testing a range of densities over several orders of magnitude - from 0.1 to 100 ng ml^−1^ per 10^5^ cells. Microbeads were coated with CD40L molecules at a maximum density, approximately 2.7 x 10^4m^ olecules μm^−2^. To achieve the maximum dose (100 ng ml^−1^ for 10^5^ cells), approximately 100 microbeads per B cell were needed. We employed two different methods to test lower doses - first by lowering the cell to bead ratio while keeping the maximum CD40L surface density constant, and second by lowering the CD40L surface density while keeping the cell to bead ratio constant (at 1 to100). B cell expansion, measured by live cell count on Day 6 of sGC culture (**Fig. 2c**), showed a clear CD40L-dose dependence. Further, the presence of soluble crosslinking antibody for soluble CD40L significantly increased cellular expansions. The expansion of B cells in sGC culture with microbead surface-bound CD40L increased in a dose dependent manner up to 10 ng ml^−1^, but decreased at the maximum dose. This decrease in B cell expansion at the maximum dose is likely due to cytotoxicity from high number of microbeads, which is supported by the fact that better B cell expansion was achieved by lowering [cell:bead] ratio to 1:10 compared to lowering the surface density of CD40L for the same dose of 10 ng ml^−1^.

Next, using flow cytometry, we investigated CSR and differentiation status of the B cells harvested at Day 6 of each sGC culture. First, the use of microbead-bound CD40L as artificial T_FH_ significantly improved the CSR, compared to the soluble systems (**Fig. 2d-f**). It was interesting to see that in the case of soluble CD40L plus soluble crosslinking antibody, the CSR to IgG1 was inversely proportional to the dose of CD40L. However, microbeads-presented CD40L not only enhanced the percentage of IgG1+ cells with increasing dose of CD40L but also induced more complete switching of isotypes indicated by an increasing shift from IgM+IgG1+ double positive to IgM-IgG1+ single positive populations and significant increased the expression of the IgG1 isotype.

Second, we also tested the differentiation status of resulting B cells by checking the expression of typical GC B cell markers, GL7+Fas+ and CD38-CD80+ (**Fig. 2g, 2h, and Supplementary Fig.1 and 2**). When we compare the two highest CD40L doses of all the culture conditions, cultures with microbead-based artificial T_FH_ cells induced equivalent (GL7+Fas+) or more (CD38-CD80+) GC B cell populations (>80%) compared to the number generated in culture conditions with soluble CD40L. Altogether, we conclude that effective recapitulation of T_FH_ cell functions was achieved by using controlled surface presentation of CD40L on the microbreads to modulate the quality and strength of CD40-CD40L ligations. The T cell dependent critical aspects of germinal center reaction, *i.e.* B cell expansion, CSR, and expression of GC B cell phenotypes, were successfully achieved *in vitro*.

It is noteworthy that this sGC reaction was also successfully applicable to B cells isolated from PBMCs (**Supplementary Fig. 3**), and completely transformed the majority of non-class switched naïve B cell populations to IgG1+ GC B cells within 5 days of culture.

**Figure 3.**
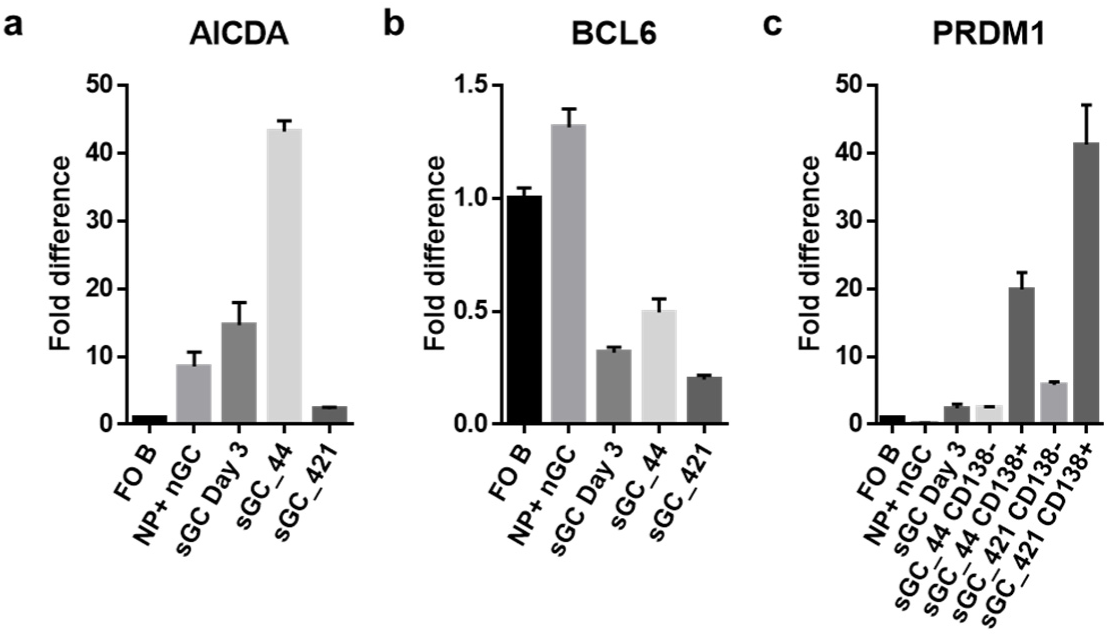
Transcriptional differentiation of sGC B cells. Quantitative real-time PCR analysis of AICDA (**a**), BCL6 (**b**), and PRDM1 (**c**) transcripts in each indicated cells. FO B: Splenic follicular B cells from unimmunized C57BL/6J mice sorted by FACS (CD19+, CD43-, CD21+, CD23^high^, CD138-), NP+ nGC: physiological (natural) GC B cells purified by FACS (NP+, CD19+, CD43-, GL7+, Fas+) from the spleen of C57BL/6J mice vaccinated with NP45-CGG /Alum 2 weeks before, sGC Day3: total sGC B cells after initial 3-day cultures performed under IL-4, sGC_44 and sGC_21: IL-4 used in the initial 3 days of sGC cultures was either maintained (sGC_44) or switched to IL-21 (sGC_421) for the rest of the sGC cultures up to Day 6. CD138+ or CD138- of sGC_44 and sGC_421 cells were also purified by FACS, accordingly. Representative data from two independent experiments are reported as the fold differences in expression levels relative to that in follicular B cells (FO B, set as 1). Error bars represent s.d. of triplicates.

### IL-4 induced sGC B cells are transcriptionally similar to centrocytes and commit to plasma cell lineage upon incubation in IL-21

We investigated the transcriptional programs involved in the sGC B cell differentiation process by using real-time quantitative PCR (RT-PCR) analysis. First, we tested the expression level of activation induced cytidine deaminase (AICDA or AID), which is known to be highly expressed in GC B cells and is involved in both CSR and SHM.^15-17^ As expected, sGC B cells express high levels of AICDA mRNA within 3 days of culture, compared to naïve follicular B cells as well as antigen-specific natural GC (nGC) B cells isolated 2 weeks after vaccination (**Fig. 3a**). BCL6 is known to be a critical transcriptional factor involved in GC reaction, as mice deficient in BCL6 are characterized by lack of GC formation and inability to produce affinity-matured antibodies.^18,19^ However, expression levels of BCL6 mRNA in sGC B cells are lower than total natural GC B cells and even lower than follicular B cells (**Fig. 3b**). This data can be explained by the fact that GC B cells are composed of two sub-stages, centroblasts and centrocytes, in early and later stages of GC reactions, respectively. BCL6 is upregulated in rapidly proliferating centroblasts and is involved in transcriptional repression of many genes to suppress the pre-mature activation and apoptosis of early GC B cells. But only a small percentage of centroblasts are further activated by T-cell dependent CD40 signaling, proceeding into centrocytes in which BCL6 expression is downregulated.^20^ The release from BCL6 transcriptional repression is required for CSR, SHM, and further differentiation of centrocytes into memory B cells and plasma cells.^21^ Therefore, the majority of our sGC B cells seem to be differentiated into centrocyte stage much more quickly than in a natural GC reactions due to the ubiquitous activation with CD40-signaling.

In a separate experiment, we observed that switching the exogenous cytokine from IL-4 to IL-21 on Day 3 of sGC culture (sGC_421) enhances the IgG1-expressing populations and the plasma cell marker, CD138-positive populations, compared to maintaining IL-4 for the whole sGC culture period of 6 days (sGC_44) (**Supplementary Fig. 4**). In order to understand these cytokine-induced phenotypic changes at a transcriptional level, we also examined the sGC_44 and sGC_421 cells by RT-PCR analysis separately. It is clear that both AICDA and BCL6 are down-regulated when IL-4 is switched to IL-21 (**Fig. 3a, 3b**). Thus, switching to IL-21 seems to slow down the IL-4 induced sGC reactions such as CSR and SHM, but further increases the speed of differentiation of GC B cells by suppressing BCL6. As it has been shown that IL-21 together with CD40L promote differentiation of centrocytes to plasma cells by upregulation of transcriptional factor Blimp-1,^22^ we also examined the expression level of PRDM1 gene that encodes Blimp-1. PRDM1 has been shown to be expressed in a small subset of centrocytes and is critically required for the formation and maintenance of plasma cells.^23,24^ Our sGC B cells at Day 3 under IL-4 already express higher levels of PRDM1 compared to follicular B cells or natural GC B cells (**Fig. 3c**). Among the sGC B cells that are maintained under IL-4 for 6 days (sGC_44), only the CD138+ cells express upregulated level of PRDM1. And upon switching IL-4 to IL-21, PRDM1 mRNA level was indeed greatly enhanced both in CD138- and CD138+ populations (**Fig. 3c**).

**Figure 4.**
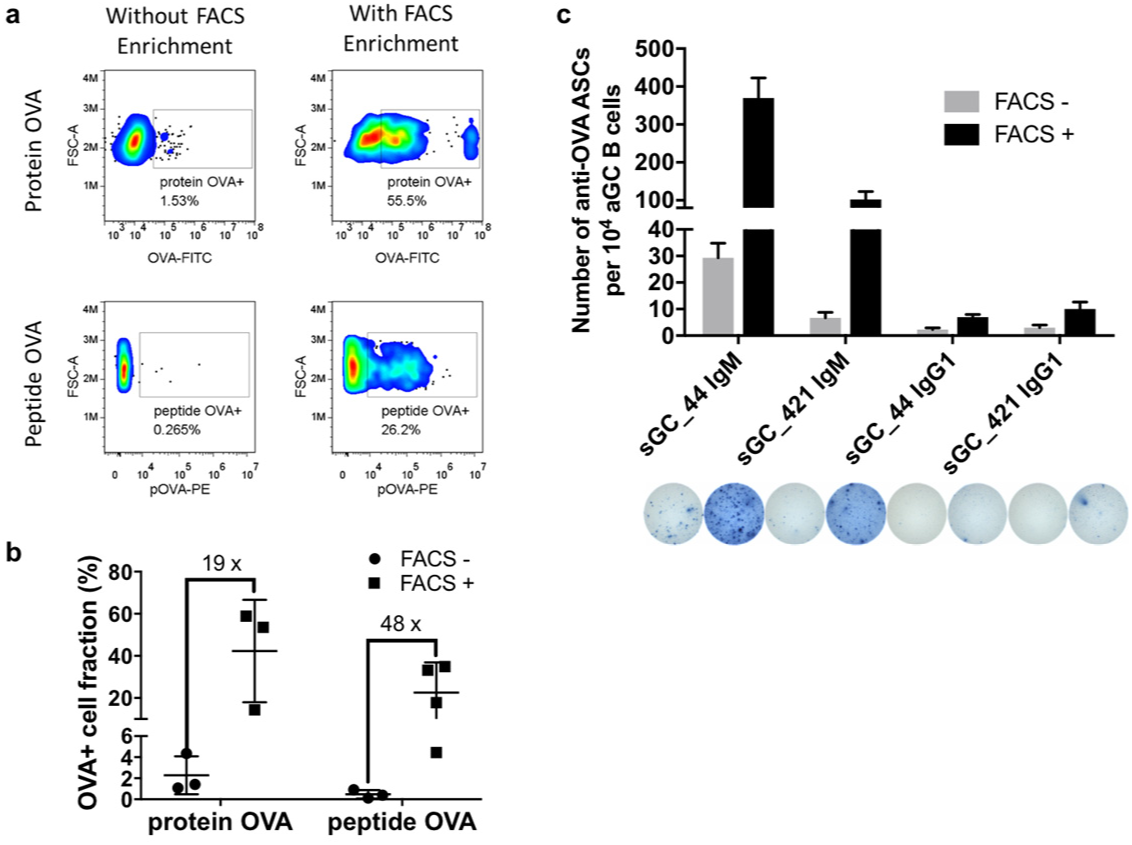
sGC reaction for antigen-specific B cells isolated by FACS enrichment. (**a, b**) Flow cytometry analysis for antigen-specificity of the resulting B cell populations after 6-day sGC cell culture. (**a**) Representative flow cytometry data for the day-6 sGC B cells stained with whole protein OVA followed by detection using FITC-conjugated anti-OVA antibody (upper panel), or stained with PE-conjugated peptide-OVA (323-339) tetramer (lower panel), with (right panel) or without (left panel) after the initial FACS enrichment step. (**b**) Percentage of antigen-specific B cell populations in total B cells after 6-day sGC culture with (FACS+) or without (FACS-) the initial FACS enrichment steps. The model antigens, protein OVA or peptide-OVA tetramer, were accordingly added into the sGC culture medium at 1 μg ml^−1^ on Day 0 and Day 3. Mean and s.d. were calculated from at least 3 separate experiments, and the indicated numbers are fold enrichments. (**c**) The frequency of anti-OVA antibody secreting cells (ASCs) among the resulting B cells after 6-day sGC culture, determined by ELISPOT assay. Shown is the mean and s.d. of triplicates. Photographs are representative images of ELISPOT wells of each condition following the order of shown in graph from left to right.

Altogether, our sGC B cells induced under IL-4 seem to be able to undergo AICDA-dependent GC reactions such as CSR and SHM, and rapidly differentiate into a centrocyte-like state by suppressing BCL6 activity. Upon switching cytokine environment from IL4- to IL-21, AICDA is down-regulated and differentiation commitment into plasma cell lineage is greatly enhanced.

### Enriched populations of functional antigen-specific B cells were effectively induced by sGC reaction following fluorescence activated cell sorting (FACS)

In order to fully recapitulate the induction of antigen-specific humoral immunity as a result of physiological GC reactions, we tested the selection of B cell populations containing antigen-specific immunoglobulin membrane receptors by use of fluorescence activated cell sorting (FACS). We first chose whole protein ovalbumin (OVA) as a model antigen, for which we tried two labeling strategies: i) labeling with unmodified OVA followed by FITC-conjugated rabbit polyclonal anti-OVA antibodies (Abcam); ii) labeling with OVA-tetramers formed between custom prepared biotinylated OVA and PE-conjugated streptavidin. The first strategy takes advantage of commercially available anti-OVA antibodies, so this method may not necessarily be applicable to every antigen. Meanwhile, it is theoretically possible to apply the second labeling strategy to any antigen of interest. However, this method carries a risk of losing some natural epitopes of a protein antigen if the amino acid within an epitope is used for the biotin-modification. Nevertheless, both strategies typically yielded ∼0.2% cells of spleen isolated naïve B cell populations that are positively stained at a varying degree of fluorescence signal with the antigen. This population is a mixture of naïve B cell clones that possess membrane immunoglobulin receptors with mixed levels of affinity toward the antigen. For the selection of a B cell population with receptors towards a more defined epitope, we used peptide-tetramers of an H-2b-restricted class II OVA epitope, OVA (323-339, ISQAVHAAHAEINEAGR). The peptide-tetramer was freshly prepared by incubation of PE-streptavidin with excess of N-terminal biotin-labeled OVA peptide followed by purification steps. Using this tetramer of a single OVA epitope, the fraction of positive staining naïve B cell population is reduced to 0.02% ∼ 0.1%, as expected.

Using these fluorescence-labeling strategies, we enriched the antigen-specific B cell populations *via* FACS before introducing them to sGC cultures. Without FACS enrichment, the staining pattern after 6 days of sGC reaction was not very different from the staining pattern of naïve B cells (**Fig. 4a**, **left**). But the FACS-enriched populations showed a significantly enhanced positive fraction in both protein (>50%) and peptide (>25%) OVA staining (**Fig. 4a, right**). On average, FACS-enriched populations contained 19 and 48 times higher fractions of antigen-specific B cells after sGC reactions compared to the non-enriched B cells for protein and peptide OVA, respectively (**Fig. 4b**). Next, we examined the number of OVA-specific antibody secreting cells (ASCs) created by sGC reactions with and without the FACS enrichment by ELISPOT assay (**Fig. 4c**). For both sGC_44 and sGC_421 conditions, the FACS enrichment produced a significantly higher number (∼14 times for IgM, ∼3 times for IgG1) of OVA specific ASCs (**Fig. 4d**), which validates the use of FACS enrichment. It is worth noting that the ASCs generated from the sGC reactions are still mostly of IgM isotype. It is similar to the physiological immune responses, where IgM is always the first class of antibody secreted from the initially generated plasmablasts. Nevertheless, by combination of flow cytometry and ELISPOT, we ultimately confirmed that the antigen-specific enrichment by FACS followed by sGC reaction will be useful for the generation of antigen-specific ASCs.

### Microbead surface-bound polyvalent antigen presentation on artificial FDCs during sGC reaction selectively increased the fraction of antigen-specific B cell population

In a physiological GC, B cells are given opportunities to survey the antigens presented by follicular dendritic cell. Those B cells that possess antigen-specific B cell receptors (BCRs) bind to, internalize, and process (degrade) the antigen, and further present the antigenic epitopes to the T cells in a major histocompatibility complex-restricted manner. This antigen-BCR interaction and cell-cycle-dependent subsequent signaling enables the clonal selection and proliferation of antigen-specific B cells by either enhancing survival of high-affinity B cells^12^ or by better antigen presentation to the T cells^13^. Thus, we tested how presenting antigens in our sGC reaction affects the antigen-specific populations. First, we compared the size of OVA-specific B cell populations after 6 days of sGC cultures with or without the presence of soluble OVA antigens. When 2 μg ml^−1^ of monomeric soluble OVA was added to sGC reactions, the OVA-specific populations grew to 1.76% compared to 0.26% without OVA during the culture (∼6 times increase, **Supplementary Fig. 5**). Using the same doses of OVA, multimeric OVA induced an expansion of OVA-specific B cells in similar quantity to the monomeric OVA.

**Figure 5.**
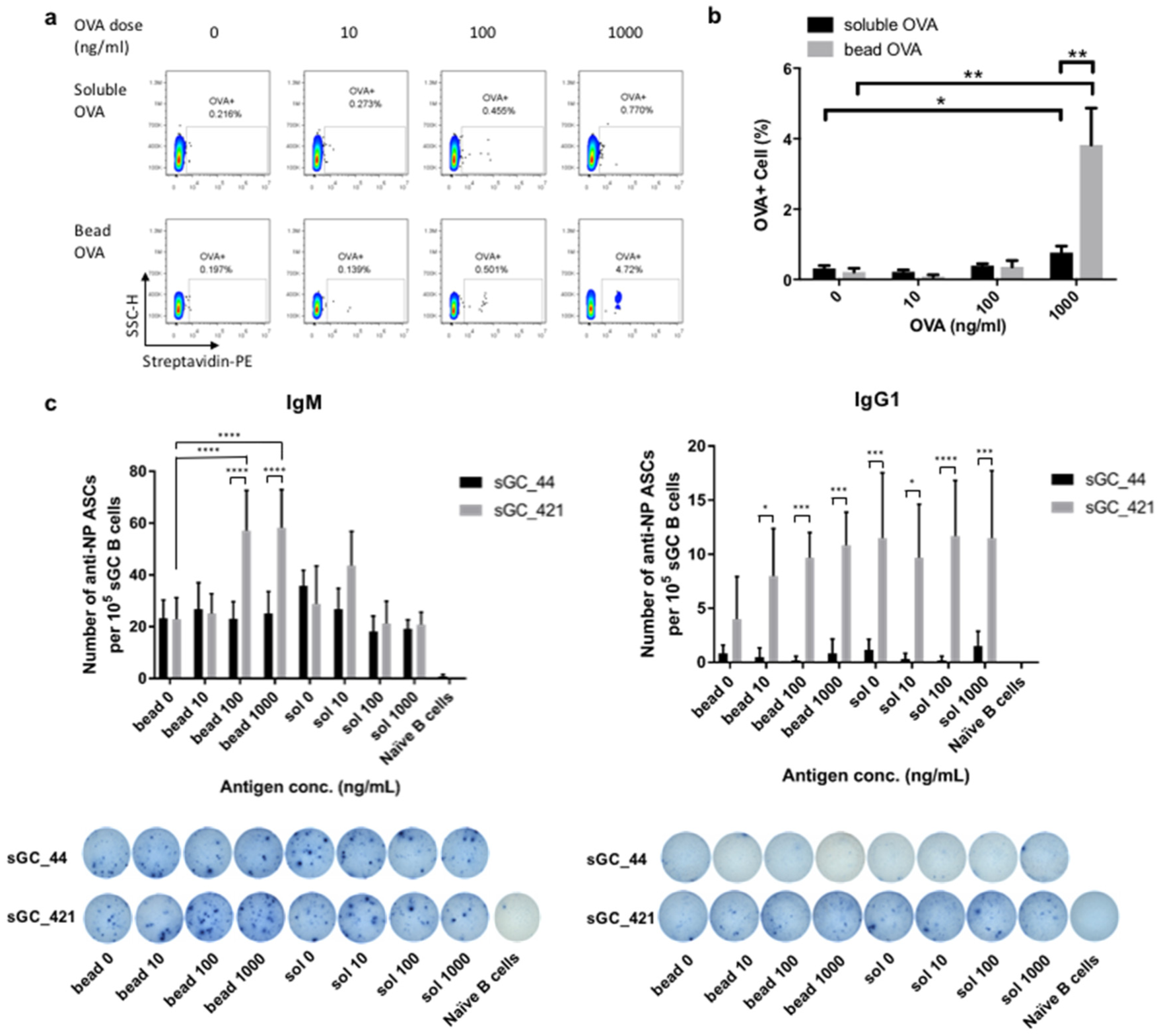
Selective increase of antigen-specific B cell populations in the presence of antigens during sGC culture. (**a**) Collection of representative flow cytometry data for resulting B cells after 6-day sGC culture stained with PE-conjugated OVA-tetramers. The indicated dose of OVA molecules was added into the culture in a soluble form (upper panel) or in a bead-bound form (lower panel). (**b**) Percentage of OVA-specific B cell populations, shown as mean and s.d. from triplicates for each condition. **(c)** The frequency of anti-NP antibody secreting cells (ASCs) among the resulting B cells after 6-day sGC culture, determined by ELISPOT assay. Shown is the mean and s.d. of n=6 replicates. Photographs are representative images of ELISPOT wells. Statistical significance for the difference between conditions was confirmed by Student *t*-test and indicated by * for *P*<0.05; ** for *P*<0.01, *** for *P*<0.001, and **** for *P*<0.0001.

Even though BCRs can directly bind to soluble antigens, B cells in *in-vivo* GC encounter a majority of antigens either as integral components of a membrane or in Fc- or complement receptor-tethered forms on the membrane of follicular dendritic cells (FDCs)^25-27^. Thus, we developed simple artificial FDCs for surface-bound presentation of OVA by incubating streptavidin-coated microbeads with biotinylated OVA molecules. The resulting artificial FDC surface presented OVA molecules in a maximum density of approximately 8.4 x 10^4^ molecules μm^−2^. Neither soluble nor bead-bound OVA brought about a significant increase in the size of OVA-specific population, up to 100 ng ml^−1^, compared to that without using OVA. However, at the maximum dose tested, 1 μg ml^−1^, soluble and bead presentation of OVA yielded 2.4 and 18 times higher OVA-specific populations compared to no-OVA-in-culture conditions, respectively (**Fig. 5a**). By direct comparison, the bead presentation created ∼5 times higher fraction (avg. 3.82%) of OVA-specific cells than the soluble presentation (avg. 0.76%) (**Fig. 5b**).

We also cultured B cells with hapten 4-hydroxy-3-nitrophenylacetyl (NP)-conjugated BSA in either soluble form or presented on an artificial FDC surface, with either IL-4 treatment for 6 days of culture (sGC_44) or combined treatment of IL-4 for 3 days followed by IL-21 for 3 days (sGC_421). While soluble NP antigen up to 1000 ng/mL did not yield a significant increase in the number of NP-specific ASCs (IgM or IgG1), bead-bound NP antigen at 100 and 1000 ng/mL resulted in an 1.85- and 1.75-fold increase of IgM+ NP specific ASCs, respectively, compared to the negative control (**Fig. 5c**). Interestingly, this increase was only observed using sGC_421 conditions for IgM, and was not observed in either sGC_44 or sGC_421 conditions for IgG1.

These data obtained for two different T-dependent antigens indicate that the antigen-BCR interactions in sGC reaction support the selective survival and/or proliferation of antigen specific B cell populations, and that surface-bound, multivalent antigen presentation induces stronger support than soluble antigens.

### sGC culture induced limited somatic hypermutation leading to potential affinity maturation

As a critical outcome of GC reactions, B cells containing high-affinity BCRs toward a specific antigen are selectively expanded to become long-lived plasma cells and memory B cells. In GCs, this affinity maturation of B cells is achieved by repeated cycles^28^ of cell division and hypermutation of IgV region genes^29,30^ under selective pressure toward higher affinity BCRs. In order to test if a similar affinity maturation can be induced during a sGC reaction, we decided to limit the model antigen to a single epitope, peptide OVA 323-339. Without the FACS enrichment, the pOVA specific population comprised only about 0.2∼0.5 % of the total B cells after sGC reaction with the concurrent use of pOVA-tetramer in culture (**Fig. 6a**).

**Figure 6.**
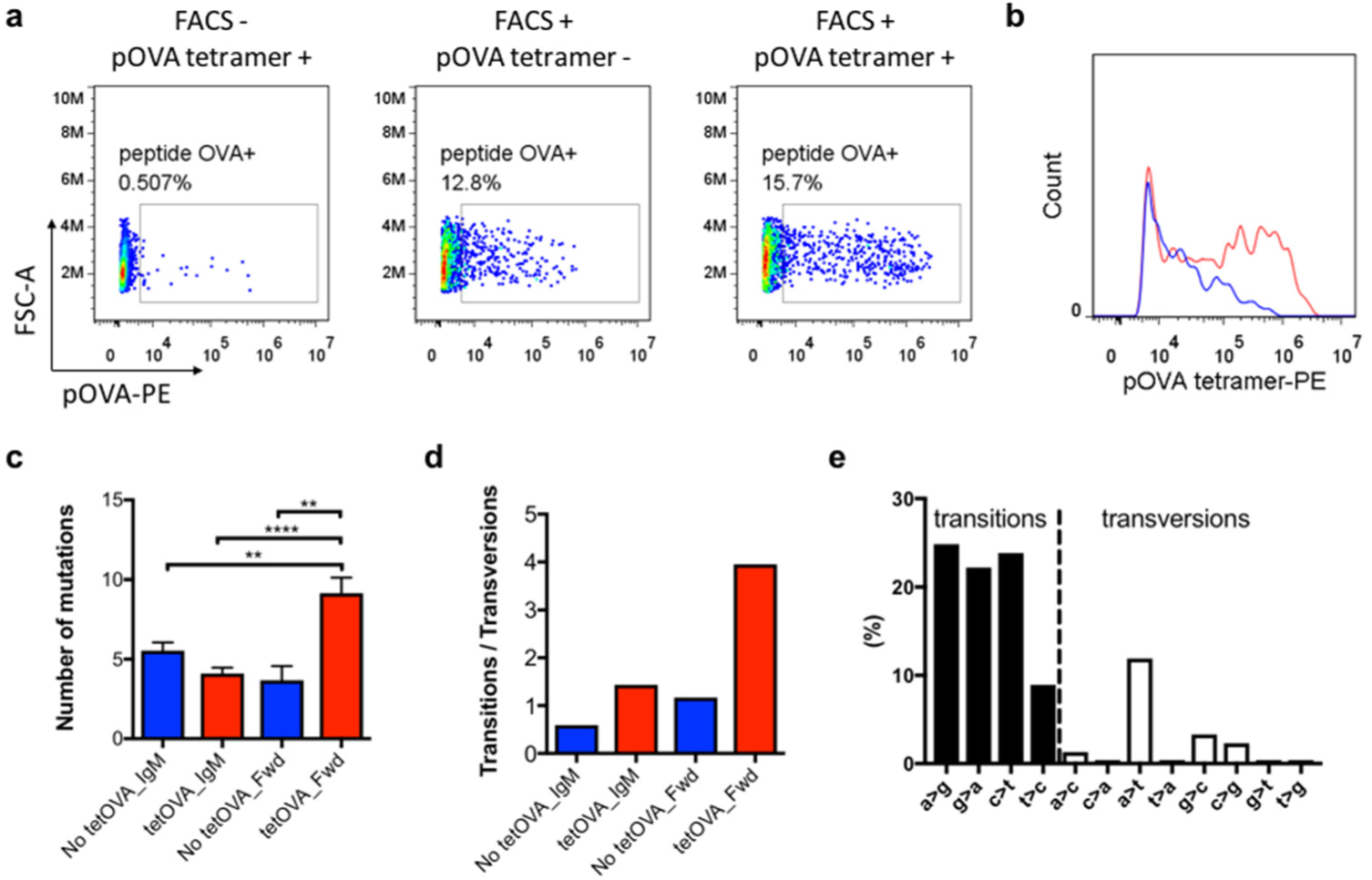
Potential affinity maturation by somatic hypermutation (SHM) in sGC reaction with multimeric epitopes. (**a, b**) Flow cytometry analysis of the B cells stained with PE-conjugated peptide-OVA (pOVA) tetramers after 6-day sGC culture. (**a**) pOVA-specific B cell populations are gated and the percentages are notified. (**b**) The histograms of pOVA tetramer-positive B cells were compared between the B cells cultured after initial FACS enrichment step (FACS+) in the presence (red) or absence (blue) of the unconjugated tetramer pOVA during the sGC cell cultures. (**c**) Number of mutations per heavy chain sequence for the pOVA-specific B cells cultured with or without the unconjugated tetramer pOVA during the sGC cell cultures. Samples contain both IgM as well as class switched IgG sequences. The values of indicated mean ± s.e.m. are 5.541 ± 0.5113, 4.104 ± 0.3695, 3.680 ± 0.8826, and 9.152 ± 0.9654, for No tetOVA_IgM, tetOVA_IgM, No tetOVA_Fwd, and tetOVA_Fwd, respectively. (**d**) The ratio of total number of transitions over transversions was calculated and graphed for each sequencing dataset. The values indicated in a bar graph are 0.593, 1.434, 1.167, and 3.951 for No tetOVA_IgM, tetOVA_IgM, No tetOVA_Fwd (forward), and tetOVA_Fwd (forward), respectively. (**e**) For tetOVA_Fwd dataset, the proportion of each indicated nucleotide switching was calculated and graphed. The filled bars represent 4 transition mutations, and the empty bars represent 8 transversion mutations. Within (**c**), statistical significance from ordinary one-way ANOVA followed by Tukey’s multiple comparisons test were represented as * (p<0.05), ** (p<0.01), and **** (p<0.0001).

To study the antigen-driven effects, we decided to enrich the pOVA specific populations by use of FACS before sGC reactions. As shown above, FACS enrichment gave rise to more than 10% of pOVA specific cells after 6 days of sGC cultures. It was intriguing that high-fluorescence staining populations emerged only from the culture with pOVA-tetramer (**Fig. 6a, b**). It is also worth noting that the similar emergence of brightly-staining populations was repeatedly observed in every sGC culture using different CD40-ligation methods, *i.e.* anti-CD40 antibody, soluble CD40L, and bead-bound CD40L, but only in the presence of tetrameric antigens during the culture.

Next, we employed a next-generation sequencing technology (454 Sequencing, Roche) to investigate the occurrence of mutations in the heavy chain of resulting BCRs from sGC reactions. For the two pOVA-specific B cell populations harvested from sGC reactions, with or without the tetrameric pOVA during the cultures, we acquired a total of 1,644 sequences, representing 6 different VH gene families and 85 clones with shared CDR3 junctions. For each population, we only analyzed productive, full-length heavy chain sequences. We analyzed the sequences acquired using forward primer (5’ VHE) separately from the sequences acquired using reverse primer (3’ Cμ). Thus the forward sequencing data represent potentially all isotype classes, while the reverse sequencing data were IgM-specific. Interestingly, when the average number of mutations per sequence were calculated for each dataset, the forward sequencing data from the sGC culture with tetrameric pOVA gave rise to a significantly high number (9.15), compared to the others (forward sequencing data without the antigen (3.68), IgM-specific sequencing data with and without the antigen, 4.10 and 5.54, respectively) (**Fig. 6c**). This indicates a potential enhancement in mutations among the isotype-switched B cells in the presence of multimeric antigens. In a further analysis to calculate the ratio of total number of transition mutations (exchange of a purine for a purine or of a pyrimidine for a pyrimidine) to transversion mutations (exchange of a purine for a pyrimidine or vice versa) in each dataset, transitions were much more favored over transversions only in the forward sequencing data acquired from the B cells cultured with tetrameric pOVA (**Fig. 6d**), among which transitions of C:G pairs slightly outnumbered those of A:T pairs (**Fig. 6e**). These data suggest that the higher number of mutations observed in this dataset is likely due to the somatic hypermutation (SHM) triggered by activation-induced deaminase (AID)^31^.

### Adoptively transferred sGC B cells engrafted in the secondary lymphoid organs, maintaining their centrocyte or memory B phenotypes

If the B cells generated from sGC reaction can be adoptively transferred, it could potentially serve as a novel autologous cell therapy regimen for the induction of long-term antigen-specific humoral immunity. In order to test the feasibility of this, we adoptively transferred approximately 1 x 10^7^ CFSE-loaded B cells from a 6-day sGC culture into non-irradiated syngeneic mice. The spleen and lymph nodes were harvested 4 days after the adoptive B cell transfer (ABCT). In flow cytometry analyses, both the spleen and the pooled lymph nodes harvested from the mice after ABCT contained higher number of CFSE-positive cells compared to the control mice without the ABCT (**Fig. 7a and b**). And these CD19+CFSE+ double positive donor B cells expressed IgG1, CD80 (B7.1), and Fas, which were not expressed in the majority of CD19+CFSE- recipient mouse B cells (**Fig. 7c**). When lymph nodes harvested 4 days after ABCT were examined using immunofluorescence microscopy, the CFSE+ cells were scattered throughout the perifollicular regions, B-cell zone as well as some in the T-cell zone (**Fig. 7d**).

In order to track the adoptively transferred sGC B cells for a longer period, we generated sGC B cells expressing genetic marker (CD45.1+) and transferred them into non-lethally irradiated (6.5 Gy) congenic host mice intravenously. After 4 weeks, donor-derived B cells (CD45.1+ CD19+) were evidently detected both in the spleen and in the lymph nodes (**Fig. 7e**). Compared to the low percentage of IgG1+ populations in the recipient B cells, about half of the donor B cells were IgG1+ (**Fig. 7e**). However, the distinctive expression levels of GC B cell makers, GL7, CD80, CD38, and Fas, which were maintained in the donor B cells for at least 4 days, have mostly disappeared by 4 weeks after ABCT to be very similar to the expression levels of naïve B cells from the recipient mice (**Fig. 7f**). These results suggest that the B cells cultured in sGC conditions can be adoptively transferred in during an autologous cell therapy to successfully engraft in the secondary lymphoid organs within short periods without losing their GC phenotype, and also survive long periods as latent B cells.

**Figure 7.**
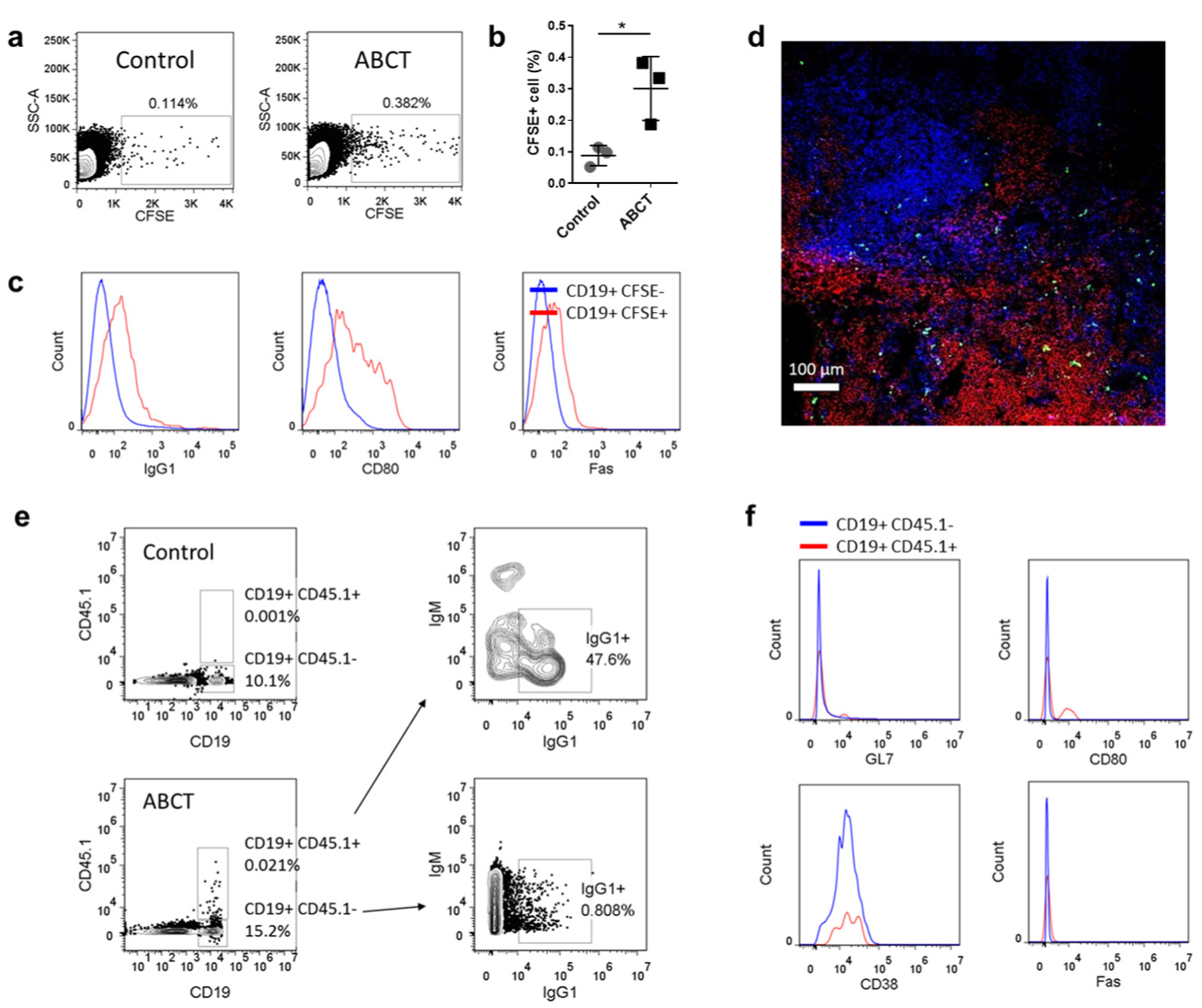
Fate of adoptively transferred B cells generated by sGC cell culture. (**a-c**) Flow cytometry analyses of the lymph nodes harvested from the recipient C57BL/6J mice 4 days after adoptive B cell transfer (ABCT) of sGC B cells derived from syngeneic donor mice and from the control mice without ABCT (saline injection). For ABCT, 1 x 10^7^ sGC B cells after 6-day sGC culture loaded with CFSE were adoptively transferred *via* tail vein. Percentage of CFSE positive cells were gated and notified from the representative flow cytometry plots for control and ABCT mice (**a**). (**b**) Collection of percentages of CFSE+ cells in the pooled lymph nodes harvested from control mice (circle) and mice 4 days after ABCT (square). Each circle and square represent a mouse. * represents statistical significance (*p*<0.05) in Student’s *t*-test. (**c**) Expression levels of the indicated markers, IgG1, CD80, and Fas, on the CD19+CFSE-(host endogenous B cells, blue) and the CD19+CFSE+ (adoptively transferred donor B cells, red) in the lymph nodes harvested from a mouse 4 days after ABCT. (**d**) Immunofluorescence micrograph of a section of an inguinal lymph node harvested from a mouse 4 days after ABCT. Green: CFSE; Red: B220 (B cells); Blue: CD4 (T cells); and the scale bar is 100 μm. (**e, f**) Representative data of flow cytometry analyses of the splenocytes isolated from the recipient C57BL/6J mice 4 weeks after ABCT and from the control mice without ABCT. Approx. 1 x 10^7^ sGC B cells derived from C56BL/6-CD45.1 mice were adoptively transferred to congenic C57BL/6J mice (CD45.2+) following non-lethal irradiation (6.5 Gy). The control mice were also non-lethally irradiated (6.5 Gy) 4 weeks before the flow cytometry. For CD19+CD45.1-(host B cells) and CD19+CD45.1+ (donor B cells) populations, relative composition of IgM+ and IgG1+ cells were enumerated. Expression levels of the other indicated markers, GL7, CD80, CD38, and Fas, on CD19+CD45.1 (host B cells, blue) and CD19+CD45.1-(donor B cells, red) populations were also compared.

## Discussion

Previously, engagement of CD40 molecules on B cells using soluble anti-CD40 or CD40L were widely used to activate B cells without comprehensive studies on their potential usage in artificial GC reactions. Here, we found that both of these approaches were only partially effective in recapitulating the GC reaction, *i.e.* the anti-CD40 antibody can induce isotype class switching in the activated B cells with minimal cell expansion and minimal expression of GC B cell phenotypes, while soluble CD40L enabled quite effective expansion and differentiation into GC-like B cells but only with minimal isotype class switching. In fact, CD40 engagement in B cells induces various biological outcomes including survival of B cells, expression and down-regulation of various surface proteins, production of numerous cytokines and chemokines, in addition to promoting germinal center formation^32^, which are achieved by activation of multiple pathways including NF-κB, MAPK, and STAT3^33^. Multiple factors mediate these complex activation pathways by direct or indirect interactions with CD40 molecules^34,35^, and it is likely that different methods of engagement for CD40 might induce an equal variety of gene expression profiles. *In vivo*, CD40L is expressed as a homotrimer on plasma membrane^2,36^, and it was shown that clustering of membrane-bound CD40L molecules is necessary for CD40-medicated B cell activation^37^. However, the 18 kDa soluble form of CD40L also naturally arises from proteolytic processing as well, all of monomeric, dimeric and trimeric forms of soluble CD40L bind to CD40 molecules on the B cell membrane and activate the B cells at a certain level.^4^,^38^

Our system of modifying microbeads with CD40L in a defined and controllable molecular density, mimicking T_H_ cells, is a rational approach to controlling the quality and quantity of CD40-mediated B cell activation. Our results demonstrate that varying doses and polyvalent presentation of CD40L on the surface of biomaterial-based artificial T_H_ cells can effectively modulate the proliferation of B cells, CSR, and the expression of GC B cell phenotypes. We have also successfully determined the optimal conditions to induce all of these canonical GC functions *ex vivo*. Together with RT-PCR analysis, we understand that this sGC reaction activates B cells to differentiate into the rapidly proliferating centroblast-like cells in the initial phase, and then quickly differentiate into the centrocyte-like B cells in which downregulation of BCL6 initiates various signaling pathways^21^. As early as 3 days following sGC culture, a significant transcriptional upregulation of AICDA, which induces CSR and SHM, is observed. At this point, a small fraction of sGC centrocytes start committing into the plasma cell lineage (CD138+) with a transcriptional upregulation of PRDM1, which is enhanced by switching the cytokine environment from IL-4 to IL-21^22^. It is worth noting that suppression of apoptosis by BCL6 is rapidly reduced in the sGC culture. Therefore, a novel culture strategy where antigen-presentation for optimal period of centroblast stage for an extended survival and expansion followed by provision of artificial T_FH_ signals for induction of centrocyte stage, or even where these stepwise reactions are alternated is worth to be explored. In the same vein, whether and how artificial presentation of other molecular signals that are present in the immunological synapse between B cells and T cells such as adhesion, co-stimulatory, and accessory receptors and ligands^39^ can be utilized for better induction and maintenance of in-vitro GC reactions remains to be seen.

In order to examine, recapitulate, and leverage the generation of antigen-specific B cells, a critical attribute of GC reactions, *ex vivo*, we used fluorochrome-conjugated multimers of whole protein model antigen OVA as well as a single B cell epitope peptide of OVA for detection and isolation of B cell populations with the corresponding BCR specificity^40^-^44^ using flow cytometry. Using our sGC reaction in conjunction with an isolation of antigen-specific B cells by FACS, we were able to routinely generate highly enriched antigen-specific GC-like B cells.

Without an extraneous enrichment step, simply including the soluble antigen in the sGC culture enhanced the fraction of antigen-specific B cell populations in a dose dependent manner. A more pronounced enhancement in the fraction of antigen-specific B cell population was achieved by providing the antigen in a surface-bound form on microbeads, which was verified with two different antigens, whole OVA protein and hapten NP. In our sGC reaction, CD40L and BAFF are provided in a nonselective manner regardless of the B cells’ affinity toward the model antigen. Thus, this observed enhancement of a selective population of antigen-specific B cells must be the additional survival and proliferative signals provided by BCR-antigen interaction, which effectively mimic the selective survival and proliferation of GC B cells upon binding of their BCRs to antigen presented by FDCs^45^,^46^. And as the concentration of antigen required for triggering effective BCR signaling is inversely proportional to the affinity toward the antigen^47^, the observed antigen-dose-dependent increase of antigen-specific B cells might be due to the BCR-dependent activation of lower affinity B cells with an increasing amount of antigen up to a certain threshold.

Moreover, it is intriguing that we repeatedly observed a B cell population that is highly stained with pOVA tetramers emerge in flow cytometry only when FACS-enriched antigen specific B cells were cultured in sGC conditions with pOVA tetramer in the culture. This brighter fluorescence could be due to either expression of higher number or BCRs or increased BCR affinity. While there is no plausible explanation for the higher expression of BCRs in these centrocyte-like B cells, there are several feasible mechanisms for the emergence of B cell populations with higher affinity BCRs. First, this could be the result of advantageous signals for survival and proliferation provided by BCR-antigen interactions to the high affinity B cell clones. Above, the same rationale was applied to account for the selective increase in antigen specific populations after sGC reactions in the presence of antigen even without the FACS enrichment step. The FACS enrichment step prior to the sGC culture simply made the selective increase in higher affinity B cells more visible. In this scenario, the presence of tetrameric epitope was essential to induce BCR-antigen affinity-dependent survival/proliferation signaling. Second, the increase in BCR affinity toward the antigen could also be introduced to the antigen-specific B cell populations as a result of somatic hypermutation (SHM) of the immunoglobulin (Ig) genes. In the GC, affinity maturation of the antibody responses is enabled by SHM of the variable regions of immunoglobulin genes^48^,^49^ and subsequent selection of higher-affinity B cell clones^50^. Similarly to CSR, SHM is known to be driven by AID (AICDA), which is largely expressed in GC B cells^15^-^17^. As previously described, RT-PCR analysis confirmed that AICDA mRNA expression is highly upregulated in our sGC culture, even higher than in the natural GC B cells. In addition, it was previously reported that SHM can be induced *in vitro* when the surface BCRs are cross-linked in the presence of help from cognate T cells^51^,^52^. From our high-throughput sequencing analyses for the V region gene of peptide-OVA-positive B cell clones, B cells emerging from sGC reactions with the addition of tetrameric peptide OVA epitope showed a significantly higher number of mutations within their potentially isotype-switched Ig heavy chains. Further, among the increased number of mutations, transition mutations were much more favored over the transversion mutations, mimicking AID-initiated physiological SHMs^31^. In our sGC model, artificial T-cell help signals are universally provided, thus the selection of higher affinity B cells cannot possibly be due to the competition in uptake of antigen for better presentation to the T cells^11^. Instead, the BCR-antigen interaction could provide not only selective signals for better proliferation and survival but also enhance the number of mutations in Ig genes, which also positively correlates to the number of cell division. Nevertheless, the majority of our sGC B cells may only be given a chance to undergo a small number of cell divisions before they are subject to apoptosis once they enter a centrocyte-like stage. Therefore, there is ample room for improvement to better support survival and increasing rounds of cell division for the antigen-selected sGC B cell populations.

Recently, cellular immunotherapies such as the adoptive transfer of antigen-specific T cells have been actively pursued for treatment of viral infections and multiple cancers including relapsed hematologic malignancies. CD40-activated B cells have been proposed either as effective antigen presenting cells (APCs) to augment T cell immunotherapies^53-55^ or themselves as effector cells in cancer adoptive immunotherapies^8^,^55^,^56^. It was encouraging to observe that the resulting B cells from our sGC reaction had migrated to and engrafted within the secondary lymphoid organs upon adoptive transfer into the non-irradiated syngeneic and non-lethally irradiated congenic host mice. And the isotype-switched donor B cells maintained Fas+ and CD80^hi^ GC B cell phenotypes for a short period (∼ 1 week), and survived for a long term (at least up to the tested period of 4 weeks) as latent B cells. Altogether, our sGC cultured antigen-specific B cells show a tremendous potential as an alternative or an adjunct immunotherapy regimen to the costly development of monoclonal antibody drugs and T cell adoptive therapies against cancers and infections^57,58^ for which no effective treatments are currently available.

## Methods

### Mice and isolation of naïve B cells

All experiments were performed using the B cells isolated from 8∼12 week old C57BL/6J or C57BL/6-CD45.1 (B6.SJL-*Ptprc*^*a*^ *Pepc*^*b*^/BoyJ) mice purchased from the Jackson Laboratory (ME, USA). All mice were maintained in the Georgia Institute of Technology animal facility under pathogen-free conditions. Procedures involving mouse were under protocols approved by the IACUC of the Georgia Institute of Technology. From the spleen, single-cell suspensions of white blood cells were prepared by simply pushing the spleen through a 40 micron cell strainer (Falcon) and incubation in ammonium chloride RBC lysis buffer (eBioscience). For the PBMC, blood was drawn by cardiac puncture from a sacrificed mouse and directly diluted in PBS with 2 mM EDTA. The PBMC were collected from the interfacial layer after a density-gradient centrifugation using Histopaque (Sigma). From these single cell suspensions, naïve B cells were further purified using a negative magnetic sorting (B cell isolation kit, Miltenyi). After depletion of cells labelled with monoclonal antibodies against CD43, CD4, and Ter-119, the resulting cell suspension typically consist of more than 97% of B cells (**Supplementary Fig. 6**).

### sGC cell culture

In a typical sGC cell culture, purified naïve B cells (10^5^ cells per well) were cultured in a 24-well tissue culture plate (Corning) in RPMI-1640 medium (Gibco) supplemented with 10% FBS (Sigma), 1 mM sodium pyruvate, 10 mM HEPES, 5.5 x 10^−5^ M β-ME, 100 units ml^−1^ penicillin, and 100 μg ml^−1^ streptomycin (Gibco). Murine recombinant IL-4 (20 ng ml^−1^, Peptrotech) and BAFF (50 ng ml^−1^, R&D Systems) were added in the initial culture medium and supplemented in every 2-3 days. For sGC_421 culture, IL-4 was replaced by murine recombinant IL-21 (10 ng ml^−1^, Peptrotech) on day 3. The CD40-CD40L ligation conditions and addition of model antigens were varied as specified in the text. For the initial comparative test for soluble CD40L vs. anti-CD40 antibody, CD40L (E61-L260, R&D Systems) or anti-CD40 antibody (clone 1C10, eBioscience) was used at 1 μg ml ^−1^. For the direct comparison between soluble CD40L and microbead-bound CD40L, recombinant murine CD40L (M112-L260) with N-terminal HA tag (YPYDVPDYA) was employed. For the crosslinking of soluble CD40L, mouse anti-HA peptide antibody (clone 543851, R&D Systems) was added at 1 μg ml^−1^. The microbeads with surface-bound CD40L molecules were prepared as artificial T_FH_ cells freshly, not more than 3 hours before initial application to sGC cell culture. Superparamagnetic microbeads (diameter of 1 μm) with covalently attached anti-HA peptide antibody (Pierce) were washed by repeating magnetic precipitation (DynaMag, Invitrogen) and resuspension in fresh PBS for at least 3 times, before incubation with CD40L molecules for 2 hours at 4 °C under rotation. For the exact control for surface density of CD40L, the binding capacity of CD40L molecules to microbeads was determined by measuring protein concentration (absorbance at 280 nm) of supernatant before and after incubation with microbeads. The dose of microbead-bound CD40L was controlled as specified in the main text for the direct comparison to the equivalent dose of soluble CD40L. In all the other subsequent sGC cultures, microbeads were employed as artificial T_FH_ cells with surface density of approximately 5 x 10^3^ μm^−2^ and maintained for the cell to beads ratio of 1:10∼1:50, unless otherwise specified. The live B cell number was counted using hemocytometer with trypan blue dye exclusion. All cultures were performed in a humidified atmosphere at 37 °C with 5 % CO_2_.

### Flow Cytometry and FACS

Cells suspended in PBS supplemented with 0.5 %(w/v) BSA and 2mM EDTA were incubated with anti-CD16/CD32 monoclonal antibody (clone 93, eBioscience) and simultaneously stained with combinations of the following antibodies: FITC-, PE-, PE-Cy7, or APC-conjugated anti-mouse CD19, IgM, IgE, IgG1, CD45R (B220), CD80 (B7.1), CD138, CD3ε, CD11b, GL7 (Ly77), CD38, and CD95 (Fas) (eBioscience). Typically 0.1 ∼ 0.5 μg of antibody was used for staining of approximately 10^5^∼10^7^ cells in 100 μl. For the control staining experiment for OVA-specific B cells, the cells were first incubated with OVA (1 μg ml^−1^), washed two times, and stained with FITC-conjugated rabbit polyclonal anti-OVA antibody (Abcam). The live cells were gated out from the dead cells and the microbeads based on FSC versus SSC (Supplementary Fig. 7). The positive gating was made based on the negative and/or isotype staining controls. All samples were analyzed using a Accuri C6 or LSR II (BD Biosciences), and fluorescence-activated cell sorting (FACS) was performed using FACSAria III cell sorter (BD Biosciences). The data were analyzed using FlowJo software. For better visualization of fluorescence signal represented on a logarithmic scale, particularly for the fluorescence units in the lower range (1-100), a biexponential transform was applied using FlowJo.

### Detection and enrichment of antigen-specific B cells

The model antigen protein OVA (Hyglos) was biotinylated using reaction with sulfo-NHS-LC-biotin (Pierce), followed by purification (Zeba spin desalting columns with 7000 MWCO, Pierce) and lyophilization. Using HABA assay (Pierce), the level of biotin incorporation was measured and the number of biotin per modified OVA molecule was determined to be 2.65. For the detection and FACS of OVA-specific B cells, multimeric OVA was freshly prepared by incubation of biotinylated OVA with PE- or APC-conjugated streptavidin in 4:1 molar ratio for 30 min at 4 °C. The incubated protein mixture was directly added into cell suspensions for the flow cytometry at 1∼10 μg ml^−1^ per 10^6^ cells. For the preparation of microbeads with surface-bound OVA molecules, streptavidin-coated superparamagnetic beads (Dynabeads with a diameter of 2.8 μm, Invitrogen) was incubated with biotinylated OVA for 1 hour at 4 °C with rotation after preparative washing steps. The loading of OVA onto the microbeads was confirmed by measuring protein concentration (absorbance at 280 nm) of supernatant before and after incubation with microbeads, and the surface density was calculated accordingly. For a model peptide epitope, N-terminal biotinylated OVA 323-339 (biotin-ISQAVHAAHAEINEAGR) was purchased from AnaSpec and used without further purification. For pOVA tetramer formation, 1 volume of 100 μM pOVA stock solution in DMSO was slowly added into 4 volume of 2 μM PE-streptavidin (eBioscience) or non-conjugated streptavidin (Pierce) in PBS and incubated for 2 hours at 4 °C with rotation, which was followed by repeated purification by filtration (Amicon Ultra, MWCO 10k, EMD Millipore). For staining of pOVA-specific population, 3∼10 nM of pOVA-tetramer was used per 10^6^ cells. For generation of NP-specific B cells, microbead-bound CD40L was provided at 50 ng/mL at a cell:bead ratio of approximately 1:50. Microbeads with surface-bound NP molecules were prepared in the same way as in the preparation of OVA-bound beads using NP_5_-BSA-Biotin (Biosearch Technologies) and provided at various cell to bead ratios (1:150, 1:15, or 1:1.5) and re-dosed every 2-3 days.

### RT-PCR analysis

RNA from each sorted B cell population was prepared with RNeasy kit (Qiagen). The cDNA was prepared by using SuperScript III First-Strand Synthesis kit and Oligo(dT)_20_ (Invitrogen). Quantitative real-time PCR was performed using RT^2^ SYBR® Green qPCR Mastermix (Qiagen) and a Step One Plus Real-Time PCR System (Applied Biosystems). Gene expression was normalized to that of β-actin (ΔCT), and a fold difference of the normalized values relative to that of follicular B cells are reported (2 to the power of –ΔΔCT). All the gene-specific primers for AICDA, BCL6, PRDM1, and β-actin were purchased from Qiagen and used without further purification steps.

### ELISPOT assays

OVA-specific or NP-specific antibody secreting cells (ASCs) generated by sGC cell cultures were detected and quantified by ELISPOT assay on a 96- well MultiScreenHTS-IP filter plate (Millipore). The plate was coated with 10 μg ml^−1^ OVA or NP_5_-BSA-Biotin for overnight at 4 °C. The cells harvested from sGC culture were washed twice before 10^4^, 10^5^, or 10^6^ cells were added to each well for at least n=3 replicates with complete RPMI-1640 cell culture media as described above. After overnight incubation in a humidified atmosphere at 37 °C with 5% CO_2_, anti-OVA IgM and anti-OVA IgG1 spots were detected by AP-conjugated goat anti-mouse IgM and IgG1 antibodies (Southern Biotech, 1:2000) in conjunction with Vector Blue AP substrate (Vector Laboratories). The image of wells were acquired using CTL-ImmunoSpot plate reader (Cellular Technology Limited).

### Immunization for generation of NP-specific natural germinal center B cells for RT-PCR

8∼12 week old mice were immunized with 100 μg of NP_45_-CGG mixed with Alum (2% Alhydrogel®, Brenntag Biosector) intraperitoneally. 2 weeks after vaccination, NP+, CD19+, CD43-, GL7+, Fas+ physiological GC B cells were isolated from the spleen via FACS.

### Next generation sequencing

The pOVA-positive B cells were sorted by FACS upon 6-day sGC cultures. The total RNA from each sample was purified using RNeasy kit (Qiagen). From the RNA, cDNA was synthesized using random hexamers and Superscript III for sequencing experiment for pOVA-specific B cells. From the cDNA, mouse Ig heavy chain V region gene transcripts were amplified by two rounds of semi-nested PCR. The PCR reaction scheme and the basic primer designs were adapted from the previous literature^59^ and slightly modified. The adapted primers originally designed by examining published Ig gene segment nucleotide sequences from the IGMT®, the international ImMunoGeneTics information system (http://www.imgt.org) and NCBI (http://www.ncbi.nlm.nih.gov/igblast/) databases were modified with the 454 Sequencing adaptor sequences and multiplex identifiers (MIDs) (**Supplementary Fig. 8**). All PCR reactions were performed in a total reaction volume of 50 μl per reaction containing 4 μl of cDNA from the previous RT-PCR as template, 200 nM each primer, 300 μM each dNTP (Invitrogen) and 1.5U HotStar Taq DNA polymerase (Qiagen). The first round of PCR was performed at 94 °C for 15 min followed by 50 cycles of 94 °C for 30 seconds, 56 °C for 30 seconds, 72 °C for 55 seconds, and final incubation at 72 °C for 10 minutes. Semi-nested second round PCR was performed with 4 μl of unpurified first round PCR product with the same reaction mixture at 94 °C for 15 minute followed by 50 cycles of 94 °C for 30 seconds, 60 °C for 30 seconds, 72 °C for 45 seconds, and final incubation at 72 °C for 10 minutes. The resulting PCR products were analyzed on 2% agarose gels, which clearly showed DNA bands corresponding to heavy chain genes with IgGs and IgM isotype. The DNA were purified from the cutout bands using Gel Extraction Kit (Qiagen). The next-generation sequencing was performed using 454 Sequencing System (Roche). All analyses for the resulting sequences were performed using IMGT/highV-Quest. The sequences showing V region identity of less than 90%, fewer than 200 base pairs, or unproductive protein translations were excluded from the analyses.

### Adoptive transfer

sGC B cells were derived by sGC_44 cultures for 5∼6 days. For a short-term tracking of the transferred syngeneic B cells, the sGC cells were loaded with CFSE (Molecular Probes) by incubation in 5 μM CFSE for 15 min at room temperature followed by washing and incubation in complete RPMI-1640 medium for 10 min at room temperature. The microbeads and dead cells were sequentially depleted from the cell suspension using a magnetic precipitation (DynaMag, Invitrogen) and negative MACS sorting by Dead Cell Removal Kit (Miltenyi) applied on LS column (Miltenyi), respectively. Approximately 1 x 10^7^ purified cells suspended in 200 ul of physiological saline per mouse were injected into mice *via* tail vein. For the short-term syngeneic or the long-term congenic transfer experiments, sGC B cells derived from C57BL6/J or C57BL6-CD45.1 mice were transferred into C57BL6/J mice (CD45.2+) with or without the non-lethal irradiation (6.5 Gy), respectively. 4 days or 4 weeks after injections, B cells isolated from pooled lymph nodes (inguinal, axillary, and brachial) and the spleen of the recipient mice and control mice were analyzed by flow cytometry and immunofluorescence microscopy.

### Immunofluorescence microscopy

The inguinal lymph nodes harvested from the recipient mice embedded in O.C.T. compound (Sakura) were frozen in liquid nitrogen and kept at - 80 °C. Frozen sections (7 μm thick) were fixed in cold acetone for 1 min. After washing 3 times with PBS, the sections were incubated with a blocking buffer (eBioscience) for 2 hours at room temperature. The sections were stained with 5 μg ml-1 anti-CD4-eFluor570 (eBioscience) and anti-CD45R-APC (eBioscience) in TBS with 1% BSA for 4 hours at 4 °C. After 3 times of rinsing with 5 minute gentle agitation in TBS containing 0.025% Triton-X and 3 times of final washing with TBS, the sections were mounted with ProLong Gold antifade medium (Molecular Probes). All the samples were examined using Zeiss LSM 700 confocal microscope (Zeiss).

### Statistical Analysis

The Student’s *t*-test was performed for verification of statistical significance. For multiple group comparison, ordinary one-way ANOVA analyses followed by Tukey’s test were performed using GraphPad Prism software.

